# MBGC: Multiple Bacteria Genome Compressor

**DOI:** 10.1101/2020.12.09.411678

**Authors:** Szymon Grabowski, Tomasz M. Kowalski

## Abstract

**Summary:** Genomes within the same species reveal large similarity, exploited by specialized multiple genome compressors. The existing algorithms and tools are however targeted at large, e.g., mammalian, genomes, and their performance on bacteria strains is mediocre. In this work, we propose MBGC, a specialized genome compressor making use of specific redundancy of bacterial genomes. Our tool is not only compression efficient, but also fast. On a collection of 168,311 bacterial genomes, totalling 587 GB, we achieve the compression ratio around the factor of 730, and the compression (resp. decompression) speed around 1070 MB/s (resp. 740 MB/s) using 8 hardware threads, on a computer with a 6-core / 12-thread CPU and a fast SSD, being about 4 times more succinct and more than an order of magnitude faster in the compression than our main competitors.

**Availability and implementation:** MBGC is freely available at github.com/kowallus/mbgc.

## 1 Introduction

Genome compression is a fairly old research topic, dating back to mid-1990s (Grumbach and Tahi, 1993). It was soon realized that even sophisticated techniques for compressing a single genome, e.g., (Cao *et al*., 2007), cannot offer much higher compression ratios than simple packing of DNA symbols into 2 bits per each. The interest of researchers thus shifted into relative compression of a genome given a reference (Christley *et al*., 2009; Pavlichin *et al*, 2013; Ochoa *et al*., 2015; Yao *et al*, 2019; Liu *et al*., 2020), typically representing the same species, or compression of a given collection of genomes without an external reference (Deorowicz and Grabowski, 2011; Wandelt and Leser, 2013; Deorowicz *et al*., 2015). We focus on the last problem variant.

The abundance of full genomes available in major repositories, like NCBI or 1KGP, in recent years poses a challenge to compress them efficiently, preferably combining high compression ratios, fast compression and decompression, and reasonable memory requirements. In this work, we focus on the compression of bacterial genomes, for which existing genome collection compressors are not appropriate from algorithmic or technical reasons.

## 2 Methods

There is significant redundancy in bacterial genomes which cannot be fully exploited using existing multiple genome compressors. The standard approach of finding repetitions between the currently processed genome and a reference genome (or possibly all previously processed genomes), and encoding them as LZ-phrases of the form (offset, length), is only moderately successful. We found out that many strings repeat as reverse-complements of corresponding strings from other genomes, a phenomenon which seemingly has not been handled earlier.

It is also beneficial not to limit the reference to one, or a few, previous genome(s), but to allow finding matches occurring almost anywhere earlier. This, however, requires a potentially unbounded memory buffer. We mitigate this problem with building the reference string, i.e., a reservoir for possible matches, in an incremental manner, appending only blocks which are “new enough”, that is, containing a relatively large fraction of DNA subsequences not seen before.

As the key ideas of our solution, Multiple Bacteria Genome Compressor (MBGC), are already sketched, now we present the algorithm in detail. The goal is to compress the sequence of genomes *G*_1_, …, *G*_*n*_ in the FASTA format. The genomes consist of one or many contigs. At the start the reference string *REF* is initialized with *G*_1_ followed by *rc*(*G*_1_), where *rc*(·) stands for the reverse complement of the passed string. MBGC also stores a literal buffer, which is initialized with *REF* (but not its reverse-complement). During the compression process, a hash table of fixed size (e.g., 2^25^ slots) is maintained, and the pairs of the form (*h, pos*) are inserted to it, where the positions *pos* are taken from *REF* accessed sparsely, with a stride of 16 symbols, and *h* are the hash values of corresponding *k*-mer seeds taken from the sampled positions. A collision on the hash *h* overwrites the previous value associated with it. In the following steps the genomes *G*_2_, … , *G*_*n*_ are taken one by one and LZ-matches of the form (*offset, length*) are sought in *REF* . The matches cannot cross contig boundaries in the current genomes. If a match is not found for the given position (note that such a check takes a constant time, due to the extremely simple hash table organization), we move to the next position in the current contig, etc., and once we have a (tentative) match, we verify its *k* symbols and try to extend it maximally in both directions. The symbols between the beginning of the current match and the end of the previous match in the current genome are added to the literal buffer. Once we are at the end of a contig, the fraction of its symbols not covered with matches is checked; if is it not small enough (exceeds 1*/*192 of the contig length, by default), the *REF* string is appended with the contig and its reverse complement. The rationale is that contigs too similar to some parts of *REF* are almost completely redundant and thus do not contribute enough to facilitate compression, but increase the memory requirement. This design decision was indeed very successful, as in our test data the string *REF* together with the concatenated literals took less than 1% of the input.

The description above corresponds to the single-threaded version of our algorithm. MBGC is, however, multi-threaded and makes use of the producer-consumer dataflow pattern. Assuming *t* worker threads, we have at most *t*−1 producers and at least one consumer for the compression. The producers decompress and parse the input (gzip) files in parallel and store them in buffers; each producer can handle up to 32 files (genomes) in its buffer. The consumer performs the actual compression (maintaining the hash table, finding LZ-matches, etc.). Once a producer fills up its buffer, it switches to compress the next unprocessed genome, which serves as a simple load balancing technique. When a genome is fully encoded, the *REF* sequence is prolonged with the relevant contigs; updates to *REF* are performed in a critical section. When the buffer of a producer is not full, the producer again fills up its buffer by reading and processing the input data, and the consumer proceeds to compress new genomes.

The resulting streams of match data (offsets, lengths), literals, header and filename data, and flags are compressed with LZMA and PPMd, using a well-known open-source software development kit (LZMA SDK).

## 3 Results

For the experiments (Table 1) we took a large collection of 168,311 bacterial genomes in the FASTA format from the NCBI Pathogen Detection project, and four 1024-genome subsets of it, each representing a single species. MBGC (in its default mode) and other compressors were tested on a Linux machine equipped with a 6-core Intel Core i7-7800X 3.5 GHz CPU, 128 GB of DDR4-RAM (CL 16, clocked at 2666 MHz) and a fast SSD (ADATA 2 TB M.2 PCIe NVMe XPG SX8200 Pro). More results and details on the datasets and the test methodology can be found in Supplementary Data.

**Table 1.** Compression results.

|  | input size<br>(GB, fasta) | HRCM |  |  | 7zip -mx=9 |  |  | zstd -19 |  |  | MBGC |  |  |
| --- | --- | --- | --- | --- | --- | --- | --- | --- | --- | --- | --- | --- | --- |
|  |  | ratio | ctime | dtime | ratio | ctime | dtime | ratio | ctime | dtime | ratio | ctime | dtime |
| C. jejuni (1k) | 1.78 | 15.0 | 224.89 | 36.45 | **61.6** | 1005.76 | 4.10 | 39.1 | 87.61 | **1.66** | 53.0 | **4.57** | 3.87 |
| E. coli (1k) | 4.87 | 158.2 | 341.29 | 44.65 | 234.9 | 1041.76 | 7.01 | 329.3 | 46.68 | 3.45 | **1302.3** | **3.33** | **1.95** |
| L. monocytogenes (1k) | 3.09 | 39.3 | 254.89 | 35.29 | 147.7 | 1105.85 | 5.24 | 94.9 | 90.09 | **2.44** | **191.5** | **3.81** | 3.02 |
| S. enterica (1k) | 5.20 | 268.0 | 340.78 | 35.30 | 248.7 | 1080.61 | 7.57 | 430.2 | 50.22 | 3.64 | **1283.0** | **3.34** | **2.00** |
| all genomes (168,311) | 587.26 | – | – | – | 173.3 | 104706.00 | 1374.17 | 181.3 | 7250.00 | 957.68 | **731.9** | **549.45** | **791.80** |

The columns “ratio” show the ratio of the input to the output size. Compress / decompress times (as “ctime” / “dtime”) given in seconds. The best results are marked bold. HRCM is single-threaded (except for the latter phase where it invokes 7zip), 7zip uses (up to) 12 threads, zstd also 12 threads and MBGC 8 threads.

As it can be seen, MBGC wins easily in the compression ratio on the *E. coli, L. monocytogenes* and *S. enterica* subsets, as well as on the total collection. It also dominates in the compression times, although not always in the decompression times. Both the compression and the decompression speed of our solution are at least on the order of hundreds of MBs per second, partly due to a multi-threaded implementation. The performance of the specialized genome compressor, HRCM (Yao *et al*, 2019), is only mediocre, and we refrained from running it on the whole collection, as the compression would take about a week. We note that some other specialized genome compressors were tried out on our data as well, with no success (for details, see Supplementary Data).

Our experiments with MBGC show that the genomes of some bacteria species can be collectively compressed by a factor exceeding 1000, at the (de)compression speed over 1 GB/s. This may be an argument for replacing the dominating gzip compression format (applied to individual genomes) with a much more resource-effective solution in DNA repositories.

## Supplementary Material to

### 1 Datasets

All the bacterial datasets used in the experiment are taken from the NCBI Pathogen Detection project (https://www.ncbi.nlm.nih.gov/pathogens) and belong to four species:

- Campylobacter jejuni https://www.ncbi.nlm.nih.gov/pathogens/isolates/#taxgroup_name%3A%22Campylobacter%20jejuni%22 (55,627 genomes, totalling 27,755 MB in gzip),
- E.coli and Shigella https://www.ncbi.nlm.nih.gov/pathogens/isolates/#taxgroup_name%3A%22E.coli%20and%20Shigella%22 (22,523 genomes, totalling 33,708 MB in gzip),
- Listeria monocytogenes https://www.ncbi.nlm.nih.gov/pathogens/isolates/#taxgroup_name%3A%22Listeria%20monocytogenes%22 (36448 genomes, totalling 32,775 MB in gzip),
- Salmonella enterica: https://www.ncbi.nlm.nih.gov/pathogens/isolates/#taxgroup_name%3A%22Salmonella%20enterica%22 (53,713 genomes, totalling 77,239 MB in gzip).

The full list of the bacterial genomes (URLs) used in our expriments is available at https://github.com/kowallus/mbgc/releases/download/v1.0/tested_samples_lists.7z

Additionally (Table 2) we use the following yeast collections (downloaded on Nov. 25, 2020):

- Saccharomyces cerevisiae ftp://ftp.sanger.ac.uk/pub/users/dmc/yeast/latest/cere_assemblies.tgz (39 genomes, totalling 486 MB in FASTA),
- Saccharomyces paradoxus ftp://ftp.sanger.ac.uk/pub/users/dmc/yeast/latest/para_assemblies.tgz (36 genomes, totalling 429 MB in FASTA).

### 2 Tested programs

The following programs, with the set parameters (e.g., for number of threads set to 12), were used in our experiments, with their results presented either in the main paper or in Section 4 of Supplementary Material.

HRCM (Hybrid Referential Compression Method) (version from 2019-Oct-12, https://github.com/haicy/HRCM/):

compression:

~~~
./hrcm compress -r <ref-file> -f <target-list-file>
~~~

decompression:

~~~
./hrcm decompress -r <ref-file> -f <target-list-file>
~~~

7-Zip (x64)

(version 16.02 from 2016-05-21, https://www.7-zip.org/):

compression:

~~~
./7z a -t7z -m0=lzma2 -mx=9 -md=1024m <archive-file> <in-file>
~~~

decompression:

~~~
./7z x -o<out-path> <archive-file>
~~~

zstd (64-bit)

(version v1.4.5 from 2020-May-22, https://facebook.github.io/zstd/):

compression:

~~~
./zstd −19 --long=29 -T12 <in-file> -o <archive-file>
~~~

decompression:

~~~
./zstd -d --long=29 <archive-file> -o <out-file>
~~~

mbgc v1.0 (https://github.com/kowallus/mbgc/):

compression:

~~~
./mbgc <target-list-file> <archive-file>
~~~

compression of mixed species collection:

~~~
./mbgc -m <target-list-file> <archive-file>
~~~

compression (max level):

~~~
./mbgc -c 3 <target-list-file> <archive-file>
~~~

compression of mixed species collection (max level):

~~~
./mbgc -c 3 -m <target-list-file> <archive-file>
~~~

decompression:

~~~
./mbgc -d <archive-file> <out-path>
~~~

### 3 Test setup

All experiments were run on a Linux (Debian) machine equipped with a 6-core Intel Core i7-7800X 3.5 GHz CPU, 128 GB of DDR4-RAM (CL 16, clocked at 2666 MHz) and a fast SSD (ADATA 2 TB M.2 PCIe NVMe XPG SX8200 Pro). MBGC is written in C++ and was compiled with gcc 10.2.0. The disk cache was flushed between runs, to have raw reads of the input files from the disk.

### 4 Additional tables and figures

Table 1 contains some results copied from the main article, but augments them with the results of MBGC in the maximum mode, and also presents four datasets containing all (rather than only 1024) genomes from particular species. Additionally, here we also show the compression (resp. decompression) peak memory usage. The MBGC compression for the whole collection (cf. the bottom row) uses a non-standard order of genomes. Namely, instead of the original order of *G*_*i*_, where *i* = 1, 2, …, *n*, we take 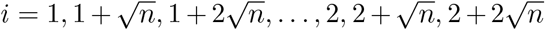 (where 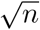 is rounded down to an integer). Such an interleaving idea helps to fill the REF sequence with contigs from various species more or less uniformly.

**Table 1.**
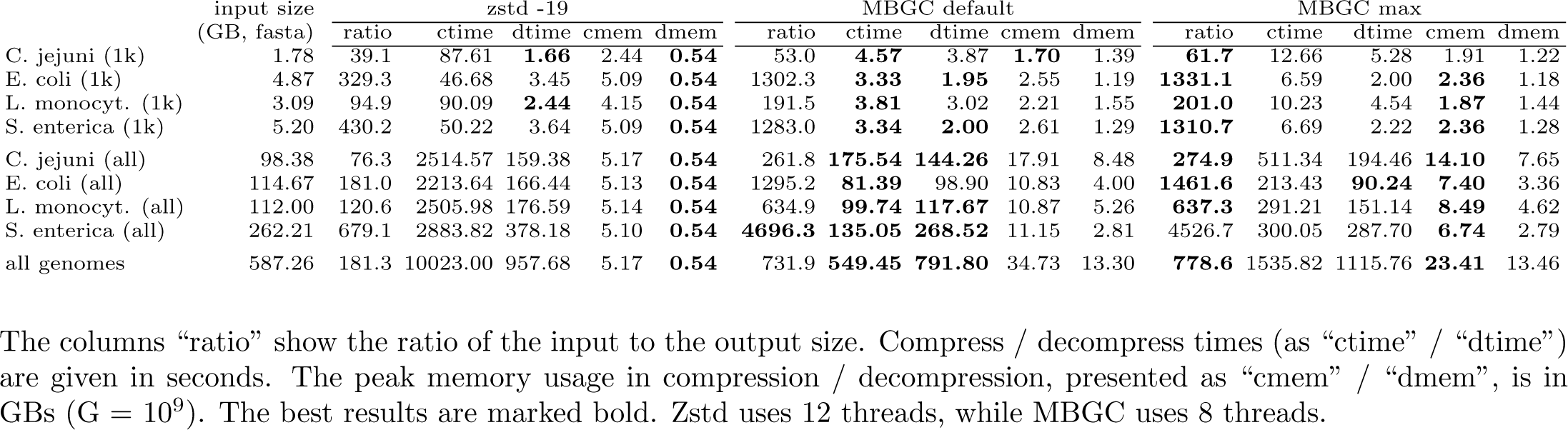
Extended compression results on bacterial genome collections.

We can notice that MBGC max (which uses changed parameters: *s* = 7, *o* = 8, and does not use threads except for parallel input and possibly gzip decompression) obtains the compression ratio by a few percent better than the default mode (with the largest gain for the *C. jejuni* data), but it is 2–3 times slower in the compression (and slightly slower in the decompression). The compression and decompression memory usage remains reasonable (although not as good as for zstd), and the max mode even tends to be more frugal than the default mode. To sum up, MBGC in the default mode is more than an order of magnitude faster than zstd −19 in compression, and marginally faster in the decompression. The gap in the compression ratio between MBGC and zstd grows with larger collections, reaching a factor of almost 7 for the whole *S. enterica*, and is about 4 for the collection of all genomes. On the other hand, zstd is more memory-frugal, which may matter if the experiments are run, e.g., on a standard laptop (MBGC default needs almost 35 GB of RAM to compress the whole collection).

The results of 7zip −mx=9 were not shown in this comparison. It needs the largest amount of memory to compress, is about 10 times (resp. 1.5 times) slower in compression (resp. decompression) than zstd and its compression ratio is a by few percent worse than zstd’s on average (although it varies for individual cases). Zstd can work with a single file input (and output) and for this reason we combined the input into a TAR archive for the zstd test (the preprocessing time for the compression process and the postprocessing time for the decompression process were not included).

We point out that End-Of-Line (EOL) symbols were removed from the DNA strings in the input files prior to the experiments, not to hamper the compression of general-purpose tools (zstd and 7zip). Preliminary experiments (with 1024-genome collections) show that on the original data (i.e., with EOLs preserved) 7zip needs about 40% more time to compress and its compression ratio is worse by a factor of 2–2.5. The respective losses are even greater for zstd (around 2.5–15 in the compression ratio, and 2–4.5 in the compression time). Such striking differences are however understandable; there are many long LZ-matches in our data, which are broken in “random” positions with the EOL characters.

We also tried rotating the input collection of 1024 genomes from the same species (by a random number from [1, 1023]), or randomly permuting it, before MGBC compression, and the compression results varied by a few and sometimes even by more than 10% in the compression ratio (the ratios for the rotated or permuted data were often, but not always, worse than with the original file order), while the compression speed was more or less proportional to the ratio, i.e., worse compression was also slower.

It may be interesting to check the impact of reverse-complement matches on the MBGC performance. It is significant indeed; according to our preliminary experiments, on *C. jejuni* and *L. monocytogenes* the compression with RC-matches turned off deteriorates roughly by a factor between 1.4 and 2.1 in the default mode.

Throughout all the presented experiments the input data are in the uncompressed (FASTA) format. Still, MBGC can read gzipped FASTA and we briefly checked how it affects overall performance. The gzipped stream is decompressed with the aid of libdeflate (https://github.com/ebiggers/libdeflate), a library for fast whole-buffer Deflate-based decompression (and compression as well, but we use it only for reading). On the individual species collections the compression time gets slightly better usually (e.g., by even around 15% for *S. enterica* and 2% worse for *C. jejuni*), with 3.5% speedup for the whole genome collection; all figures with respect to the default mode of MBGC. The compression ratio varies a little (due to unpredictable access to genomes with the worker threads), usually below 1%.

As it is often the case in bioinformatics, dealing with very large data, the I/O speed has a significant impact on the overall performance. To this end, we note that *cached* read of the data (i.e., without flushing the disk buffers, which is standardly performed in our tests) makes MBGC default about 1.5 times faster in the compression, assuming that input is not much larger than the available RAM memory. It means that replacing our SSD with an (even) faster one would also have a similar effect. This observation corresponds to the uncompressed FASTA input; we have not tried to measure such an effect with gzipped input.

We have to admit we tried to compare our software also with other tools for genome compression: GDC 2, iDoComp and memRGC. The experiments were not successful though. GDC 2 refused to compress due to uneven number of contigs in the input files. memRGC hanged during the compression. Regarding iDoComp, we experimented with *C. jejuni* (1024-genome collection). It could not process two of our genomes (a problem with header parsing), so we removed them from the input. The compression time was about 90 times longer than from MBGC default (408.1s vs 4.57s) and the resulting archive was by a factor of 8.7 times larger. We found those results not satisfactory (in other words, iDoComp is inappropriate for this kind of data) and performed no more tests with it.

Fig. 1 presents the impact of varying four MBGC parameters (*u, s, k, o*) on the overall compression ratios and compression times, on four bacterial species; each dataset is limited to 1024 genomes. We present these parameters in the next paragraphs. In the subfigures, only one parameter is modified at a time, while the others have default values.

**Figure 1:**
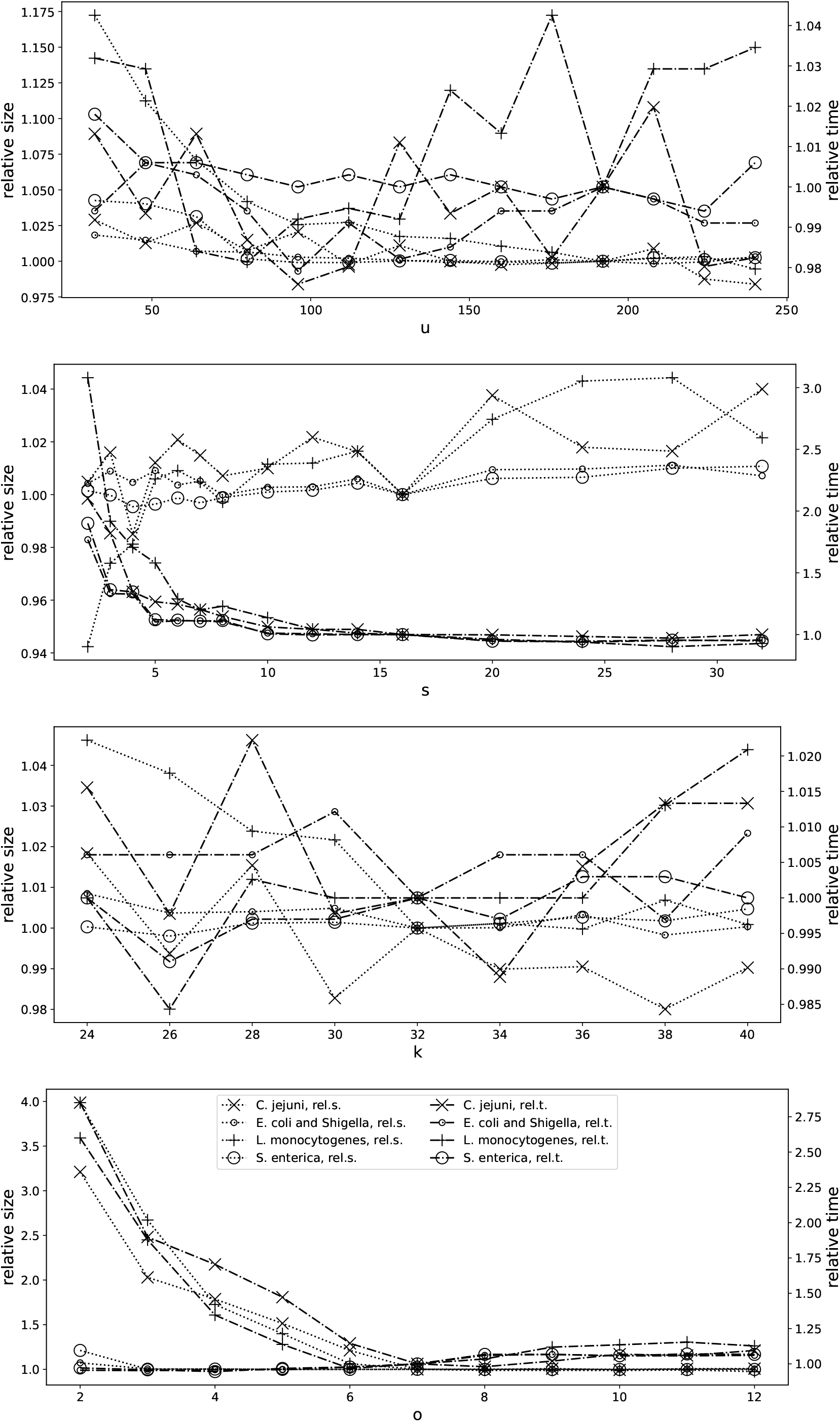
Relative compression ratios and times when one of the parameters is varied and the remaining three parameters keep their default value. The left (resp. right) Y axes are related to relative compressed sizes (resp. compression times).

The parameter *u*, or rather its reciprocal, determines the “growth rate” of the *REF* string. Its default value, *u* = 192, means that if and only if the fraction of the symbols in a just processed contig which is not covered with matches exceeds 1*/*192, then the *REF* string is appended with the contig and its reverse complement. Smaller values of *u* imply fewer contigs to meet this criterion, which makes the resulting *REF* shorter (which, however, does not necessarily reduce the compression memory usage, as a shorter *REF* tends to produce more literals, stored in the memory to be compressed later).

The parameter *s*, with the default value of 16, stands for the sampling step over the reference string. Larger *s* means that fewer substrings from *REF* become seeds to be inserted in the hash table (HT), which in turn makes the overall HT update faster and also requiring less memory (which is however also related to the parameter *o*, described later). On the other hand, with larger values of *s* some tentative LZ-matches may be missed.

The parameter *k*, whose default value is 32, is the minimum match length. It must be not less than the seed length (which is 28 by default). A too small value of *k*, which thus also implies short seeds, leads to many collisions (where new entries overwrite the old ones in HT), while a too large *k* may disallow some short matches, and the hashing itself is more costly (although this is hardly an issue, if *k* is within reasonable limits).

Finally, the parameter *o*, which stands for “referenceFactorBinaryOrder” in the source code, sets an upper bound on the *REF* length, as 2^*o*^(|*G*_1_| · 2), where |*G*_1_| is the length (in bytes) of the first genome in the processed collection, and the factor 2 corresponds to storing both *G*_1_ and its reverse complement. For example, if *o* = 7 (the default value) and assuming that |*G*_1_| is exactly 5 MB, we obtain that *REF* is limited to 1280 MB in its length. This parameter also affects the HT size. We assume the number of its slots is at least 2^24^, but in fact its number of slots is 2^*j*^ for such maximal *j* that 2^*j*^ *<* |*REF*_*max*_ |*/s*, where *REF*_*max*_ is the just discussed limit of the *REF* length, *s* is the sampling step, and *j* is additionally upper-bounded by 31. The rationale for such a HT size is quite obvious, as we do not want to have more inserts to HT than its number of slots (as repeating seeds, for reasonable values of *k* and *u*, are not that frequent).

Let us now comment the results. The first observation is that we cannot speed up the default mode of MBGC more than just a little varying one of the presented parameters. On the other hand, we can it make much slower (by a factor 2 or more) in the compression by setting small values to *s* and/or *o*, and only a small value of *s* of the options considered here may give a small to moderate compression gain; a small value of *o* is a clearly bad choice. The impact of the parameter *k* on both the compression ratio and speed is fairly small in the whole presented range of values, from 24 to 40. The compression is also fairly stable with varying *u*, if only it is not too small (note a compression loss exceeding 10% for the *C. jejuni* dataset when *u* is around 50 and below).

Fig. 2 shows the compression ratio and compression speed with varying the number of threads from 1 to 12. The speed does not improve with more than 6 threads (but perhaps it would if the compression were performed in the RAM memory). The compression ratio is rather unaffected for *E. coli* and *S. enterica*, but using already more than 1 thread for *C. jejuni* and *L. monocytogenes* yields a few percent compression loss. For *C. jejuni* the gap is as large as about 12% when the number of threads grows from 1 to (the default) 8. On the other hand, using 8 threads is about 3–4 times faster than 1 thread in the compression for all four datasets and for this reason we find the compression loss in half of the cases rather acceptable.

**Figure 2:**
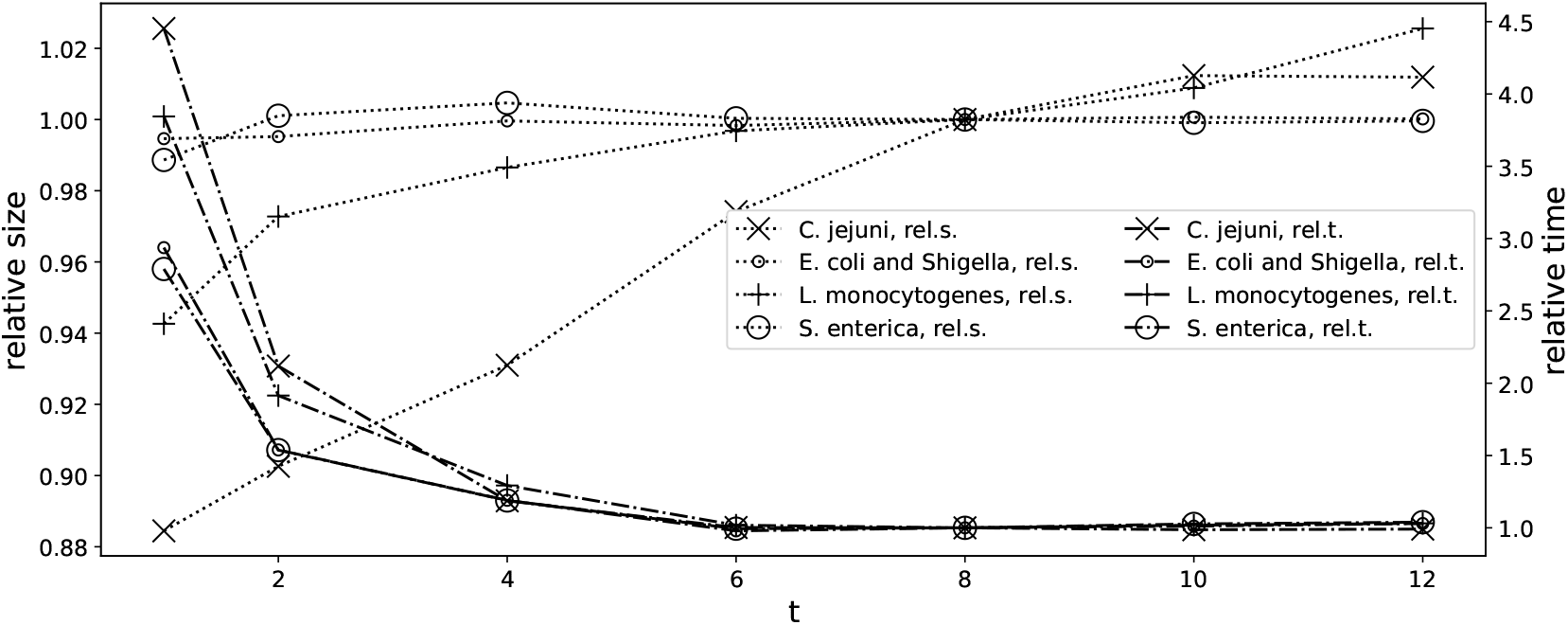
Relative compression ratios and times in the function of the number of threads. The left (resp. right) Y axes are related to relative compressed ratios (resp. compression times).

Finally, in Fig. 3 we can see how the compression ratio and compression times change when more and more genomes are given as the input. As expected, the compression time grows roughly linearly (note the X-axis scale), but the compression ratio improves, as for further genomes more similar “pieces” can be found in the already processed collection (or, to be more precise, in the currently used *REF* sequence). The only exception is *E. coli*, where after processing about 2,000 genomes the compression ratio first deteriorates somewhat and then no longer improves.

**Figure 3:**
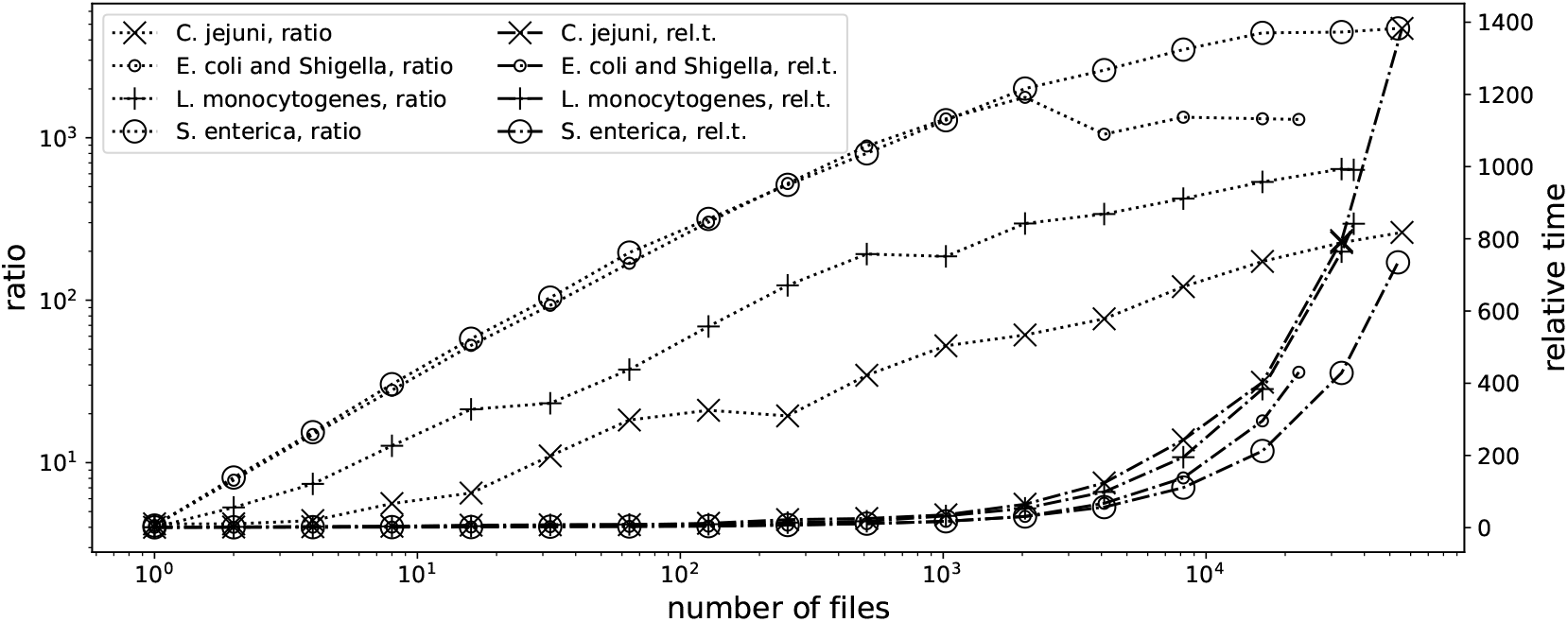
Compression ratios and relative times when the number of input genomes grows. The left (resp. right) Y axes are related to compressed ratios (resp. relative compression times).

For a separate experiment, we took two non-bacterial genome collections, *S. cerevisiae* and *S. paradoxus* (Table 2). We didn’t expect MBGC to be competitive here, and indeed, GDC 2 and 7z are superior in the compression ratio (while HRCM is on a par with MBGC max), but MBGC remains the fastest tool in the compression process. A better overall choice is, however, GDC 2, with a significantly higher compression ratio and slower by only 40% in the compression with respect to MBGC max on *S. cerevisiae*. On the other hand, the compression speed difference is more than 8-fold (in favor of MBGC max) in case of *S. paradoxus*. In decompression, zstd is the fastest (yet it handles single files, e.g., in the TAR format), followed by GDC 2 and 7z (which was also tested with a single file), and then by MGDC (default, then max). HRCM is the slowest in the decompression, but relatively fast in the compression.

**Table 2.** Compression results on non-bacterial genome collections, *S. cerevisiae* and *S. paradoxus*

|  | *S. cerevisiae* |  |  |  |  | *S. paradoxus* |  |  |  |  |
| --- | --- | --- | --- | --- | --- | --- | --- | --- | --- | --- |
|  | ratio | ctime | dtime | cmem | dmem | ratio | ctime | dtime | cmem | dmem |
| HRCM | 78.8 | 8.85 | 3.61 | 1.18 | **0.06** | 52.6 | 10.01 | 4.11 | 1.18 | **0.07** |
| GDC 2 | **109.8** | 4.97 | 0.63 | **0.52** | 0.14 | 80.7 | 28.04 | 0.99 | **0.51** | 0.17 |
| 7z -mx=9 | 100.6 | 455.70 | 0.97 | 4.92 | 0.49 | **83.8** | 430.76 | 0.91 | 4.41 | 0.44 |
| zstd -19 | 75.5 | 46.76 | **0.49** | 0.70 | 0.49 | 41.9 | 47.40 | **0.47** | 0.65 | 0.43 |
| MBGC default | 75.8 | **1.74** | 1.01 | 1.95 | 0.79 | 44.9 | **2.17** | 1.27 | 1.87 | 0.75 |
| MBGC max | 78.9 | 3.54 | 1.20 | 2.54 | 0.73 | 56.9 | 3.31 | 1.42 | 2.52 | 0.66 |

